# Identifying (un)controllable dynamical behavior in complex networks

**DOI:** 10.1101/236323

**Authors:** Jordan C Rozum, Réka Albert

## Abstract

We present a technique applicable in any dynamical framework to identify control-robust subsets of an interacting system. These robust subsystems, which we call stable modules, are characterized by constraints on the variables that make up the subsystem. They are robust in the sense that if the defining constraints are satisfied at a given time, they remain satisfied for all later times, regardless of what happens in the rest of the system, and can only be broken if the constrained variables are externally manipulated. We identify stable modules as graph structures in an *expanded network*, which represents causal links between variable constraints. A stable module represents a system “decision point”, or trap subspace. Using the expanded network, small stable modules can be composed sequentially to form larger stable modules that describe dynamics on the system level. Collections of large, mutually exclusive stable modules describe the system’s repertoire of long-term behaviors. We implement this technique in a broad class of dynamical systems and illustrate its practical utility via examples and algorithmic analysis of two published biological network models. In the segment polarity gene network of *Drosophila melanogaster*, we obtain a state-space visualization that reproduces by novel means the four possible cell fates and predicts the outcome of cell transplant experiments. In the T-cell signaling network, we identify six signaling elements that determine the high-signal response and show that control of an element connected to them cannot disrupt this response.

**Author summary:** We show how to uncover the causal relationships between qualitative statements about the values of variables in ODE systems. We then show how these relationships can be used to identify subsystem behaviors that are robust to outside interventions. This informs potential system control strategies (e.g., in identifying drug targets). Typical analytical properties of biomolecular systems render them particularly amenable to our techniques. Furthermore, due to their often high dimension and large uncertainties, our results are particularly useful in biomolecular systems. We apply our methods to two quantitative biological models: the segment polarity gene network of *Drosophila melanogaster* and the T-cell signal transduction network.

## Introduction

A key goal in the study of complex dynamical systems is to extract important qualitative information from models of varying specificity (e.g., [1, 2]). This has been approached via the construction and analysis of qualitative models (e.g., discrete models [3-7]) and also by analytic techniques applied to continuous systems [8-13]. In this work, we present and implement a new approach to identifying control-robust subsystem behavior that can drive the dynamics of the system as a whole. Our approach applies to a large class of continuous, discontinuous, and discrete models.

Interacting systems are partially described by their regulatory networks. In these networks, nodes represent each of the various interacting entities within the system, and signed edges represent direct positive or negative influence. To better understand the temporal character of the system, one can construct a dynamical model on the regulatory network. First order Ordinary Differential Equations (ODEs) are a natural choice for such models. The influence upon the value of each entity, *x_i_*, is represented as *ẋ*_*i*_ = *F_i_* (***x***), where the dependence of *F_i_* upon *x_j_* is consistent with the influence of entity *j* upon entity *i*. A validated model can be used to gain practical insights about the system, such as how to drive it into a desired attractor.

There are two key challenges to the construction and analysis of ODE models of complex interacting systems. First, there is often large uncertainty in measurements of variable and parameter values. Second, these systems are typically high-dimensional, which complicates phase-portrait visualization and other traditional qualitative analyses.

One approach to these challenges is to choose a more qualitative model. Discrete models have been used to successfully model many biological phenomena, including pattern formation and multistability [3, 4]. Despite the vast reduction in state-space afforded by discretization of variable values, exhaustive searches for dynamical behaviors are computationally infeasible in high-dimensional systems. Several methods for identifying the causal structure of state-space in discrete models have been proposed, including hierarchical transition graphs [14] and prime implicant graphs [15].

An analogous concept in ODE models is that of positive invariant sets (also called “trap spaces”) [16, 17]. These are regions of state space that system trajectories may enter but not exit. By identifying such spaces, one may make predictions about the evolution of a system without integrating the governing ODEs.

A second strategy is that of examining features in the dynamical repertoire that arise directly from the associated regulatory network and weak assumptions about the form of the dynamic model. Structural controllability relates branching patterns in the regulatory network to the identification of control targets that are sufficient to drive linear dynamics on the network into any state [18]. This allows one to study system control near a steady state. In many biological and chemical systems, however, the dynamics are nonlinear and large disruptions from equilibrium are of interest. In such systems, even when the dynamics are not specifically known, regulatory feedback loops provide useful information for global control [19-23]. For example, given relatively permissive continuity and boundedness assumptions, an ODE-described system can be driven into any of its attractors by controlling any set of variables whose removal eliminates all feedback loops and external inputs [19, 21, 24]. Positive feedback loops in particular are associated with the presence of multistability [8-12], which has been of particular interest in biomolecular systems because it is necessary for cell-fate branching and decision making [4, 25-28].

Two existing approaches to identifying the effects of positive feedback loops are especially relevant here. The first of these is the methods put forth by Angeli and Sontag for studying monotone input-output systems (MIOS) [29]. Their approach identifies steady states and their stability in systems lacking negative feedback loops or incoherent feed-forward loops (in the general meaning of two directed paths of opposite sign between two nodes). The second is based on the concept of *stable motifs* of Boolean dynamical systems [22, 30]. This method constructs an auxiliary network that encodes the regulatory logic within its graph structure (in a similar vein as logic hypergraphs ([5, 31]), enabling efficient identification of the system’s dynamical repertoire. Within this auxiliary network, certain graph structures, called stable motifs, correspond to positive feedback subsystems that sustain steady states that are impervious to influence from the rest of the network (see S1 Appendix section 1 or [30] for further details). In other words, stable motifs determine positive invariant sets. This observation connects the concept of positive invariant sets to the regulatory network in the Boolean case. Our work extends this connection to the continuous case.

Our framework encodes the causal relationships between variable constraints as the network structure of an *expanded network*. An edge from one constraint (e.g., *x >* 0) to another (e.g., *y >* 0) indicates that the first (*x >* 0) is sufficient to maintain the second (*y >* 0). The expanded network helps to identify low-dimensional subsystems that drive higher-dimensional dynamics. We show that *stable modules*, source-free expanded subnetworks subject to certain consistency criteria, correspond to control-robust positive-invariant sets of the originating dynamical system. Variables obeying stable module constraints must be directly controlled (i.e., either receive exogenous input or be made control variables) if the constraints are to be broken. This identifies variables that must be controlled to disrupt certain behaviors (or, equivalently, it identifies variables that cannot be controlled in such a way as to disrupt the behavior).

It is non-trivial to choose relevant variable constraints for the modeled system, but in practice, the form of the regulatory functions often suggests natural candidates. Furthermore, we leverage MIOS techniques to algorithmically specify meaningful constraints in a class of systems common in biology (see S1 Appendix section 2). This is implemented (S1 Source Code) as code that systematically scans for stable modules in an input ODE system satisfying certain assumptions. Identifying several stable modules in a systematic search highlights “decision points” in subsystems that determine system-wide outcomes.

## Results

### Stable Modules Describe Control-Robust Behavior

The core of our analysis strategy lies in the interpretation of an auxiliary network that is constructed from the dynamical system of interest. Following previous work in Boolean systems [3, 32], we call this auxiliary network an *expanded network*. An expanded network must be constructed from a given dynamical system. It is a network on a node set consisting of statements about the values of variables (or, equivalently, consisting of the regions of state-space in which these statements are true). There are two types of directed edges between nodes. The first type, the *maintenance edge*, indicates that one statement cannot become false while the other is true. The second type, the *driving edge*, indicates that the sustained truth of the first statement implies that the second statement will eventually become true. In this paper, our focus is on continuous, autonomous ODE systems, although the concepts are presented in such a way as to be readily adapted to other types of dynamical systems. In the following, we describe the nodes and edges of an expanded network in more detail. In S1 Appendix section 3, we provide a formal mathematical foundation for the following discussion.

In an expanded network, there are two types of nodes: virtual and composite. Virtual nodes are statements about the values of dynamic variables that can be assigned a definite truth value at any given time (e.g., the virtual node “*x >* 0” is true only when the value of the variable *x* is positive). Virtual node statements can be viewed as regions of state-space, and are true at time *t* if *x* (*t*) is in the corresponding region. Composite nodes also take Boolean values, and correspond to the composition of virtual nodes by “AND” (Λ) rules. Each composite node receives directed edges from its factor virtual nodes. As such, all factors of a composite node must be represented as virtual nodes in the expanded network. For example, the composite node *x >* 0 Λ *y >* 0 is true only when *x* and *y* are positive, and there are directed edges from *x >* 0 and *y >* 0 to this composite node. In deterministic finite-level systems, it is possible to choose a finite number of statements that fully characterize the state space [33], but in general, the nodes of an expanded network embody partial information about the system. For a given choice of virtual and composite nodes, the expanded network is unique, however, a different choice of virtual nodes for the same system can lead to different expanded networks. Some choices of virtual nodes are therefore more illuminating than others, and choosing an informative set of virtual nodes is not always straightforward. In the next section, we propose and implement a method to address this difficulty in a particular class of systems. The remainder of this section covers general expanded network properties, which are prerequisite for the methods of the next section.

Virtual nodes can receive two types of edges: a maintenance edge or a driving edge, with the latter being a more restrictive version of the former. If a virtual or composite node *X* must be false before a virtual node *Y* can change from true to false, we say that *X maintains Y* and we draw a directed edge from *X* to *Y* in the expanded network. Note that if a virtual node *X* describes a positive-invariant set in state-space that remains positively invariant even under control of variables not involved in the definition of *X*, then according to this definition *X* maintains *X*, which results in a self-loop on *X*. This can happen if *X* describes a self-activating variable, for example. In determining whether *X* maintains *Y*, we must consider all valid variable values that might disrupt *Y* when *X* is true. These variable values are drawn from the region of state-space in which the model is valid and experimentally accessible, e.g. a box in state-space defined by the maximum and minimum values of each variable. By considering values from this region of validity we simultaneously evaluate a large number of system trajectories and control strategies. To explore whether control of one system element can drive the system as a whole into particular regions of state-space, one may also wish to impose the condition that an edge from *X* to *Y* indicates that the truth of *X* implies the truth of *Y* in finite time (or, more briefly, *X drives Y*); this additional constraint is unnecessary when considering self-sustaining behavior.

A subnetwork, *S*, of an expanded network, *N*, is a *stable module* if it satisfies three conditions: (i) all nodes *X* in *S* have a parent (regulator) node in *S* (possibly *X* itself if it has a self-loop), (ii) if a composite node 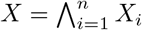 *X_i_* is in *S*, then *X_i_* is also in *S* for *i* = 1*..n*, and (iii) it is possible for all nodes in *S* to be simultaneously true. For brevity, we refer to subnetworks satisfying conditions (i), (ii), or (iii), as *source-free*, *composite-closed*, or *consistent*, respectively. Our key result is the following: if all nodes in a stable module are simultaneously true (i.e., if in that instant the system is in a region of state-space for which all virtual node statements in the stable module are true), then they remain true under all state-space configurations in the region of validity. In the following we will call a stable module whose nodes are simultaneously true an *active* stable module.

To prove our key result, consider by way of contradiction an active stable module, *S* that deactivates. Let *Y* ∈ *S* be a virtual node that becomes false before or concurrently with any other node in *S*. Because every node in *S* has a parent node in *S*, there is *X* ∈ *S* that maintains *Y*. By the definition of maintenance edges, *X* (or one of its factors if it is composite) must become false before *Y* does, violating the selection criteria and thereby proving the result.

A stable module with no stable submodules is a *stable motif*. Under the condition that a stable module, *S*, is active, we can simplify the expanded network by removing any edges that point from a virtual node in *S* (e.g., *x >* 0) to a composite node outside of *S* (e.g., *x >* 0 Λ *y >* 0) because the condition expressed by this edge is now satisfied.

We can also remove any node that is necessarily false when *S* is active (e.g., if *S* contains the node *x >* 0, the node *x <* 0 can be removed). Stable motifs of the modified expanded network are then identified and added to *S* in the original expanded network. We thus iteratively form larger stable modules, building a sequence of stabilized subsystems that drive system dynamics. When the activity of a stable module in one sequence implies the inactivity of at least one stable module in another sequence, these sequences are mutually exclusive. Collections of mutually exclusive sequences describe the system’s dynamical repertoire.

Our definition of stable motifs encompasses the definitions of stable motifs given in [30] for Boolean systems (see S1 Appendix section 1) and in [33] for multi-level systems. This allows us to generalize many results from discrete modeling to general dynamical systems. In particular, generalizing arguments in [22], we consider system control via expanded network topology. It is often of interest to identify variables that can activate a stable module (which may correspond, e.g., to a healthy cell state). This can be achieved by solving the graph-theoretic problem of identifying stable module *driver nodes*. A module *driver node set D* of module *M* in an expanded network is a set of virtual nodes *D* such that the truth of all nodes in *D* implies the truth of all nodes in *M* in finite time. Therefore, identification of a driver node set for a stable module prescribes a control strategy to trigger the module behavior. Conversely, if a stable module represents undesired behavior (e.g., a diseased cell state), one might seek to disrupt it. Because stable modules are self-sustaining, control of variables not represented in the undesired module can never achieve this goal. Disruption of a stable module requires direct control of at least one of its represented variables.

To illustrate the method, and some of its utility, we analyze a toy example (Fig 1, Eq. 1). In this toy example, we will choose statements for virtual nodes somewhat arbitrarily, with the goal of illustrating how relationships between nodes in the expanded network can be identified and interpreted. In later sections, we introduce a more systematic approach to selecting virtual nodes that does not rely on the intuition of the investigator.

**Fig 1.**
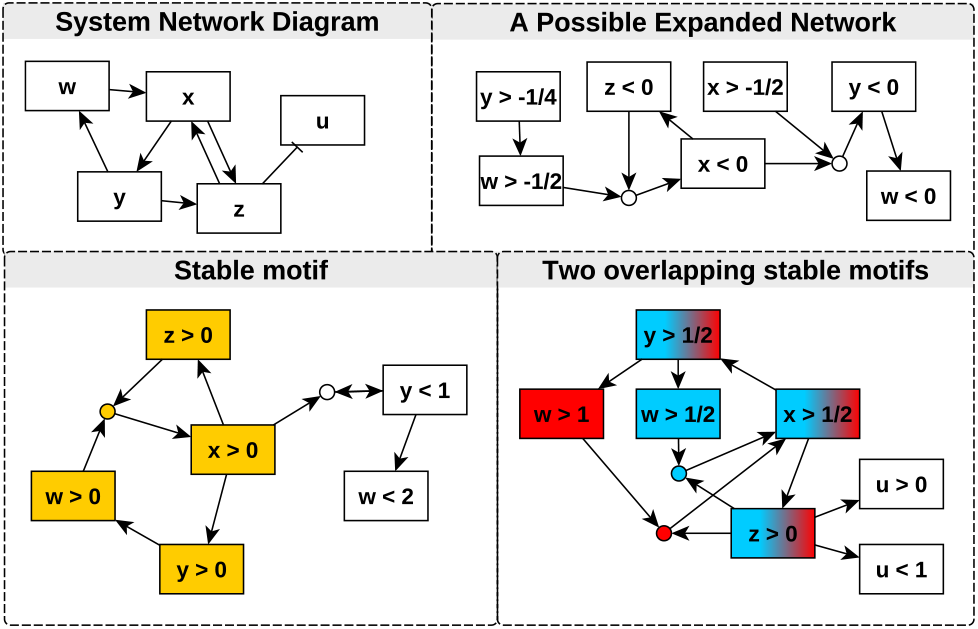
A network representation of the system given in Eq. 1 along with three expanded networks. Each circular node represents a composite node formed by composition of its parent nodes by an “AND” rule. Highlighted components are stable motifs (and therefore also stable modules). These represent conditions that, once satisfied, remain satisfied. In the overlapping motifs (marked in blue and red), we may choose to consider the motif containing *w >* 1 (red), which gives more information about the value of *w* when the motif is realized, or we may consider the *w >* 1*/*2 (blue) motif, which is more readily realized (i.e., the threshold is smaller). By considering both motifs together, we see that the *w >* 1*/*2 (blue) motif drives the *w >* 1 (red) motif, i.e., states satisfying *w, x, y >* 1*/*2 and *z >* 0 will eventually also satisfy *w >* 1. We remark also that the stable motifs shown could be expanded to incorporate the other nodes depicted in the components. Such structures are stable modules and are also self-sustaining, but include nodes that are not necessarily part of any feedback loop; they are instead driven by feedback elsewhere in the network.

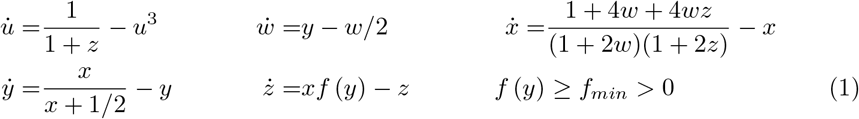

Here, we have very limited information about *f* (*y*); perhaps it is stochastic or discontinuous. Nevertheless, by uncovering upper and lower bounds on components of the ODE vector field, we can begin to assemble an expanded network one edge at a time. For instance, if *x >* 1*/*2 holds, then *ż> f_min_/*2 *− z* is implied. If *z* is positive and decreasing (*ż <* 0), it cannot decrease faster than *f_min_/*2 *− z*. In this case z would asymptotically approach *f_min_/*2. As a consequence, *z* will never fall to zero. Therefore, as long as *x* is greater than 1*/*2, *z* cannot fall below 0 once it has become positive, and so we say that *x >* 1*/*2 maintains *z >* 0. A similar argument applies in any case when *x* is larger than an arbitrary positive value. Furthermore, if *z* is not positive, then *ż* is strictly greater than *f_min_/*2. Therefore *z* will eventually (in finite time) become larger than zero and so we say *x >* 1*/*2 drives *z >* 0. We can therefore conclude that there is an edge from *x >* 1*/*2 to *z >* 0 in the expanded network. Similarly, we see that *x* will be maintained above 1*/*2 if *w >* 1*/*2 and *z >* 0 are both true. We therefore identify a composite node (*w >* 1*/*2) Λ (*z >* 0) with incoming edges from *w >* 1*/*2 and *z >* 0, and an outgoing edge to *x >* 1*/*2. We continue to identify edges in the expanded network and search for stable modules. Some of the subgraphs of the expanded network that can be generated in this way are depicted in Fig 1 alongside the traditional network representation of the system. We have identified three stable modules, thereby proving, for example, that if the systems satisfies *x, y, w, z >* 0 at any time, it will always satisfy those conditions (as follows from the yellow module in the bottom left of Fig. 1). The other two modules contain *x, y >* 1*/*2 and *z >* 0 as well as either *w >* 1 or *w >* 1*/*2 (see bottom right panel of 1). Thus, if the system satisfies the four conditions given by either module, it will continue to do so for all time. The arguments underlying the construction of the subgraphs of the expanded network hold for any *f* (*y*) *> f_min_ >* 0, and so we have extracted meaningful qualitative information despite large dynamical uncertainty. In addition to the expanded networks and their subgraphs containing stable modules, many that do not contain stable modules also exist (e.g., the top right panel of 1). Such networks contain information regarding the consequences of directly controlling particular nodes so that they satisfy virtual node statements (e.g., if we fix *y <* 0, we see that *w* will eventually become negative).

Choosing virtual nodes defined by inequalities, as is our main focus here, has important implications for how oscillations are observed. If a variable oscillates, but remains above or below some threshold, the statement indicating the variable value relative to that threshold can be part of a stable module. Alternatively, oscillations can manifest in the expanded network as subnetworks with contradictory virtual nodes. For instance, if *ẋ* = *z* + *sin*(*y*) *− x*, then *z > z*_0_ (where *z*_0_ is an arbitrary positive number) maintains (and drives) *x > z*_0_ − 1 and *z < z*_0_ maintains (and drives) *x < z*_0_ + 1.

The main difficulty in identifying stable modules is determining what statements are most useful for inclusion in the expanded network. If the statements are too general, then either the results will not provide much insight, or the network will be too sparse because the statements are not sufficiently restrictive to imply one another. If a statement is too restrictive, on the other hand, it may have an in-degree of zero in the expanded network, in which case it cannot be part of a stable motif. Despite these challenges, we have found a straightforward approach to analyzing threshold behavior of a large class of biologically relevant systems.

### Application to Biological Systems

We consider a broad class of dynamical systems that take the form

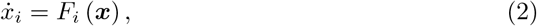

where *F_i_* is continuous, monotonic in each of its arguments, and strictly decreasing in *x_i_*. This class of ODEs describes many biological systems (S1 Appendix section 2) and is particularly well-suited to analysis in our framework.

The essential steps of the stable module identification process are as follows. First, we identify all subgraphs of the regulatory network that are composed of positive feedback loops. For each such subgraph, we construct two families of candidate stable modules by conjecturing that each variable *x_i_* in the regulatory subnetwork has a virtual node of the form *x_i_ > T_i_* or *x_i_ < T_i_*, where *T_i_* is left unspecified (for brevity, we denote this form by 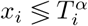). For each candidate module, we construct a “worst-case” monotone system by replacing any variable regulatory effects that would introduce a negative feedback loop or incoherent feedforward loop by constant regulatory effects. This system is analyzed using the techniques of [29] such that equilibria of the worst-case system yield thresholds *T_i_* for which the candidate stable module is genuine. In the following we provide the details of the process.

To each variable *x_i_* of a regulatory subnetwork under consideration, we assign a set of thresholds 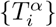 and consider virtual node statements of the form 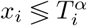. At this stage, each 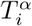 may remain parameterized and the statements need not cover the full dynamical range of *x_i_*. We create composite nodes 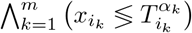 as needed. Next, we conjecture that particular edges exist in the expanded network for some (unspecified) choice of threshold parameters. For instance, when activity of one variable, *x*_1_, is sufficient for activation of another, *x*_2_, we would hypothesize the formation of an edge *x*_1_ *> T*_1_ →*x*_2_ *> T*_2_. In the conjectured expanded network, we find source-free, consistent, and composite-closed subgraphs, which serve as candidate stable modules.

Consider a candidate stable module, *S_c_*, in the conjectured expanded network. Consider also the regulatory subnetwork, *G_c_*, made up of nodes represented in *S_c_* and all incident edges. Some of these incident edges are represented in *S_c_*, while other “external” edges are not. For example, a candidate stable module *S_c_* in Fig 1 might be *y > T_y_ → w > T_w_ → x > T_x_ → y > T_y_* and the corresponding regulatory subnetwork *G_c_* consists of the positive cycle x, y, w and the additional external edge from z to x. We note that external edges may exist between two nodes in *G_c_* if the regulatory relationship between these variables is not part of *S_c_*. To identify bounds for the virtual nodes that ensure that the candidate stable module *S_c_* is genuine, we use the monotone input-output systems (MIOS) methods of Angeli and Sontag [29], which apply to sign-consistent systems (see S1 Appendix section 4).

The relationships represented in *S_c_* constitute a sign-consistent subnetwork (S1 Appendix section 4). Any sign-inconsistencies in *G_c_* arise from external edges. To construct a sign-consistent modified subsystem for *S_c_*, we consider each variable *x_i_* represented in *S_c_*. Any external regulation of *x_i_* by *y_j_* is held fixed by replacing *y_j_* with a “worst-case” value in *F_i_*. The “worst-case” value is chosen such that *x_i_* is as close as possible to 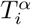 in the stable module node 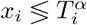; because *F_i_* is monotonic in each argument by assumption, this is either *y_j_ ≡* inf *y_j_* or *y_j_ ≡* sup *y_j_* (i.e., when *y_j_* is as large or small as possible within the region of validity). For example, if *y_j_* negatively regulates *x_i_* and 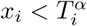 is in *S_c_* then we evaluate *F_i_*|_yj=inf yj_.

Because the resulting modified subsytem is sign-consistent, we can apply the MIOS procedure of Angeli and Sontag (Theorem 3 of [29]). For examples of this process for sign-consistent systems, see [20, 29]. To do this, we must verify that we can select a variable, *x_k_*, called the “MIOS input variable” that has the property that maintaining *x_k_* at a constant value drives the system to a single steady state for all initial values of variable other than {*x_k_*} [20, 29]. The form of Eq. 2 implies that a node in the modified system satisfies these conditions if its removal makes the modified system acyclic.

Once we have verified that a MIOS input variable can be chosen, we can follow Angeli and Sontag [20, 29] to find the steady states of the modified subsystem. These steady states determine the thresholds that we use for the virtual nodes in *S_c_*. The sign-consistency of the modified subsystem implies that these thresholds describe a positive invariant set of that subsystem ([29]). This sign-consistency together with the monotonicity of the regulatory functions implies that this set remains positively invariant for all possible values of the external regulatory effects because any deviation in these from their worst case values unambiguously drives the system away from the boundary of the stable module subspace and into its interior. Therefore, with these thresholds, *S_c_* is realized as a valid stable module for the original system.

We illustrate this method by identifying a candidate module and constructing a worst case system in the example of Eq 1. First, we recall that we have already shown that the system is restricted to the positive orthant if the initial conditions are within this region, so we assume that this is our region of validity. In general, identification of the region of validity often follows from physical or biological considerations. By inspection, we observe that *y* activates *w*, which activates *x*, which in turn activates *y*. We thus conjecture that a stable module of the form *y > T_y_ → w > T_w_ → x > T_x_ → y > T_y_* exists. Note that this feedback loop is positive and defines a loop closure of a monotone system when *z* is viewed as a parameter. To identify valid bounds for this candidate stable module (if such bounds exist), we construct the worst case system for the candidate. As the only regulatory effect not represented in the candidate is the effect of *z* on *x*, we must identify the value of *z*, within the region of validity, for which *x*̇ is minimized. In this case, *x* ̇ is minimized when *z* is maximized, and so we allow *z* to tend toward infinity in the worst case system, yielding a worst case system given by 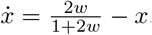, along with *ẇ* and *ẏ* from Eq 1. The steady state of this system is given by the solution of the feedback characteristic equation 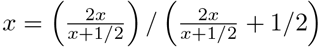, which has solution *x* = 7*/*10, yielding *w* = 7*/*6 and *y* = 7*/*12. We thus conclude that *y >* 7*/*10 →*w >* 7*/*6 →*x >* 7*/*10 is a stable module. We provide additional examples in sections 5 and 6 of S1 Appendix.

We have algorithmically implemented (S1 Source Code) this process by considering intersecting unions of positive feedback loops. For each union, we conjecture two stable modules (in which one set of nodes is “high” and the other is “low”, and vice versa). Using user-specified physical system bounds, we construct a “worst case system” for each candidate stable module, as described above, and test the existence of a MIOS input variable. If such a variable can be found, we use it to numerically find the steady states via the MIOS procedure. If any steady states are within the physical system bounds, we return the corresponding stable module.

The above procedure returns a list of stable modules involving threshold statements about subsystem variables connected by positive feedback loops. Note that generally the list of stable modules we generate does not directly correspond to all of the system’s equilibria, or even necessarily to equilibria at all. Rather, it corresponds to “trap” subspaces, i.e., positive-invariant sets, that are robust to control of regulatory effects external to the subsystem. If the control includes multiple regulatory effects, we assume that these effects can be controlled independently of each other. The list of stable modules generated for each subsystem is in one-to-one correspondence with the equilibria of this subsystem that are robust to such control. This list thus contains the subsystem behaviors that are self-sustaining under all control strategies that preserve the topological structure of the regulatory network. Additional behaviors may be robust to only a subset of these interventions.

In the remainder of this paper, we use the above methodology and automation scheme to analyze two systems from the literature. The first, the Drosophila segment polarity gene network, is a prototypical system used to study a broad class of embryonic pattern formation mechanisms. The second example is the T-cell signaling network, which is a characteristic representative of signal transduction networks, which lead to specific cell responses to environmental signals.

### Single-Cell Drosophila Segment Polarity Network

The original multicellular model of the Drosophila segment polarity gene network [34] uses coupled ODEs to model the concentrations of mRNAs and proteins of a family of genes that are important for the development of segments in *Drosophila melanogaster* embryos (see Fig 2). This family of genes includes engrailed and cubitus interruptus, which encode transcription factors, as well as wingless and hedgehog, whose proteins are secreted and interact with proteins in the neighboring cells [34-36]. We use a modified version of this model (equations 12-23 in [35]), which has incorporated more recent experimental results (e.g., on the *sloppy-paired* protein) and been recast for a single cell while assuming steady-state values for neighboring cells. Because no measured values of the kinetic parameters in the model are available, and because our purpose here is illustrative, we have simply chosen parameter values from the biologically relevant parameter region (see S1 Appendix section 7 for parameter values and variable abbreviations).

**Fig 2.**
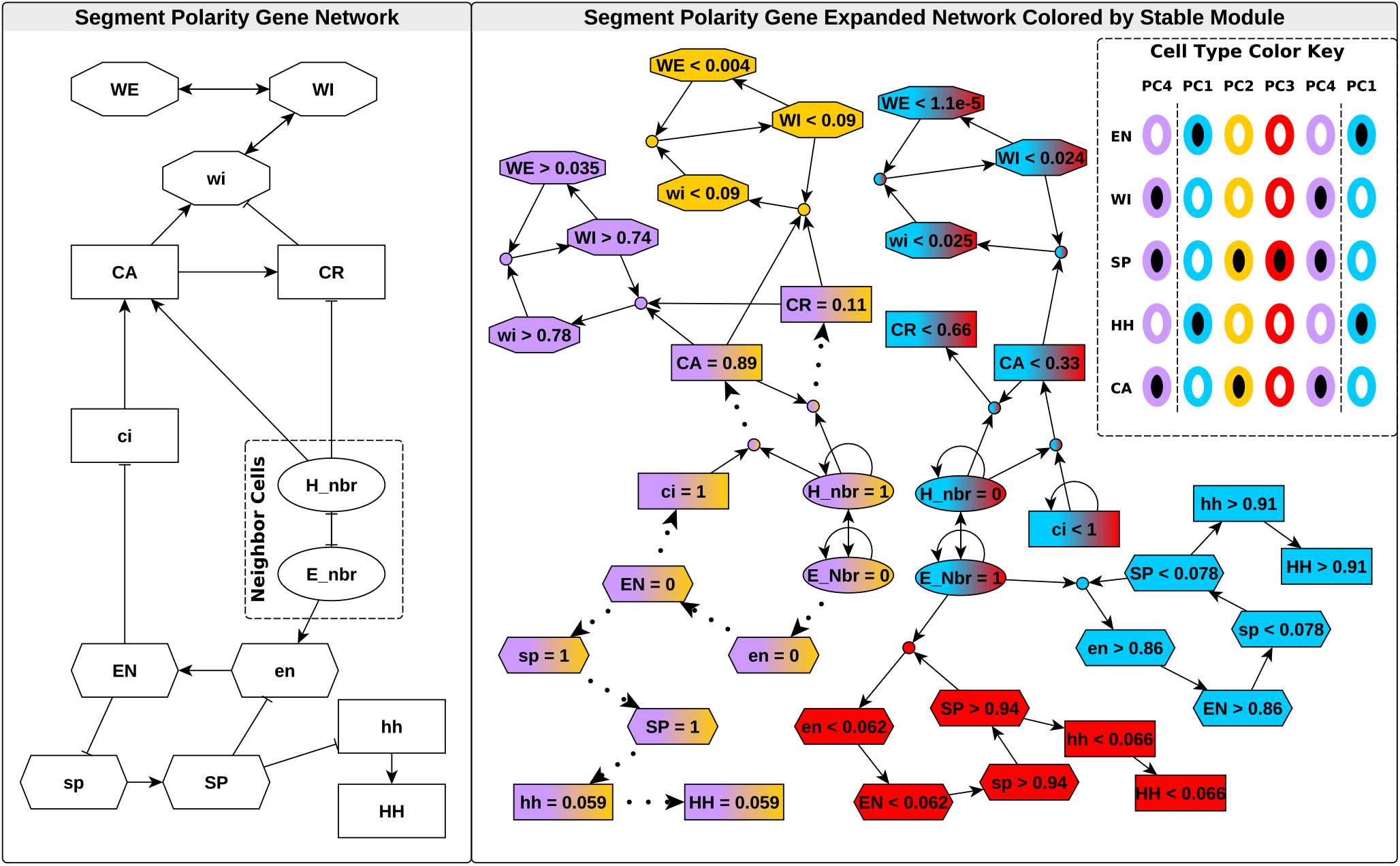
The network schematic for the single-cell model of *Drosophila* segment polarity genes [35] (left) and expanded network (right). The color key (right inset) summarizes observed characteristics of cells in the *Drosophila* embryonic segments (parasegments) ([34-36]; see S1 Appendix section 7 for parameter values and the full names of abbreviated variables). Each column represents an individual cell, arranged by anterior-posterior position in the parasegment. Columns are colored and named according to cell type, which is determined by prevalence of the proteins labeling each row. Black-filled ovals represent high levels of the protein, while white-filled ovals represent low levels. In the expanded network (right), dotted lines represent the node maintenance relation and asymptotic implication. Solid lines indicate the maintenance relation and implication in finite time. Nodes are colored according to membership in each of four biologically relevant stable modules, which correspond to the four cell types identified in the right inset. Node shape indicates participation in feedback loops that sustain these stable modules.

We identify several stable modules of biological importance in this model. When neighboring cells have high levels of *wingless* protein, we find two stable modules distinguished by differential *sloppy-paired* and *engrailed* expression (red and blue nodes in Fig 2). For high concentrations of neighboring *hedgehog* protein, we find two stable modules involving the *wingless* sub-network (yellow and purple nodes in Fig 2).

By shading the nodes in the expanded network according to module membership (as in Fig 2) we can visually identify regions of state-space that correspond to different attractors of the system. Specifically, these attractors distinguish the four cell-types observed in the development of *Drosophila melanogaster* segments, which we label PC1-PC4 [34, 36] (see Fig 2). Furthermore, the expanded network highlights the causal chains that link regions of state-space and establish cell fates. By identifying driver node sets for stable modules, we can prescribe control strategies to attain any of the four cell types. For example, drivers of the cell type PC1 (blue module in Fig 2) are high neighboring *hedgehog* (*H_nbr_*) and low *sloppy-paired* (*sp* or *SP*).

Furthermore, we can use this information to form hypotheses about the outcome of altering node states. For example, we can make the following prediction about the outcome of a future wet-bench experiment in which a cell of a certain type is transplanted to a region in which neighboring cells express *hedgehog* and *wingless* at higher or lower levels relative to the cell’s initial neighbors. Consider a cell of type PC1 (blue module in Fig 2). If the neighboring *wingless* (*E_nbr_*) and *hedgehog* (*H_nbr_*) are reversed in expression level, that disrupts the *engrailed* -*sloppy paired* part of the module. As a result, *en* and *sp* approach zero and one, respectively. The values of *wingless* (*wi*) and the two configurations of its protein before transplant are consistent with the stable module characterizing cell type PC2 (yellow module in Fig 2). Therefore, our analysis of the model ([35]) suggests that a qualitative change in cell gene expression from that of the foremost cell of the embryonic segment (PC1, blue module) to that of the second segmental cell (PC2, yellow module) would be observed in such a transplant experiment. Numerical simulations support this conclusion (S1 Figure). Our analysis also identifies the reason for this change: the *engrailed* -*sloppy paired* feedback loop is not robust to elimination of neighboring *wingless* (*E_nbr_*). If this prediction is falsified by follow-up experiments, the lack of transition would imply the existence of additional regulation of *engrailed* and/or *sloppy paired*. The additional regulation would need to act in such a way as to allow a high expression of *engrailed* in the absence of neighboring *wingless*.

### T-Cell Receptor Signaling Network

The second biological example we consider here is a model that describes the cascading activation of transcription factors when T-cell receptors are bound by external molecules [37]. The model was constructed using the *Odefy* MATLAB toolbox ([38]) to transform a pre-existing Boolean model of T-cell activation ([31]) into an ODE model *ẋ*_*i*_ = *F_i_* (***x***) = (*R_i_* (***x***) *− x_i_*) */τ_i_*, where each *R_i_* is a polynomial of Hill functions with *R_i_* (***x***) ∈ [0, 1] describing the regulatory effects that influence the production of *x_i_*. The parameters *τ_i_* are the inverse degradation rates of the various biomolecules.

To simplify the example, we consider the strongly connected core of the system with saturated input signals, though the precise signal strength has little impact on the analysis. The resulting network is depicted in Fig 3 (left), in which the edges are labeled with the Hill coefficient, *n*, and disassociation constant, *k*, of the function *H_i_* (*x_i_*) for the corresponding regulatory effect.

**Fig 3.**
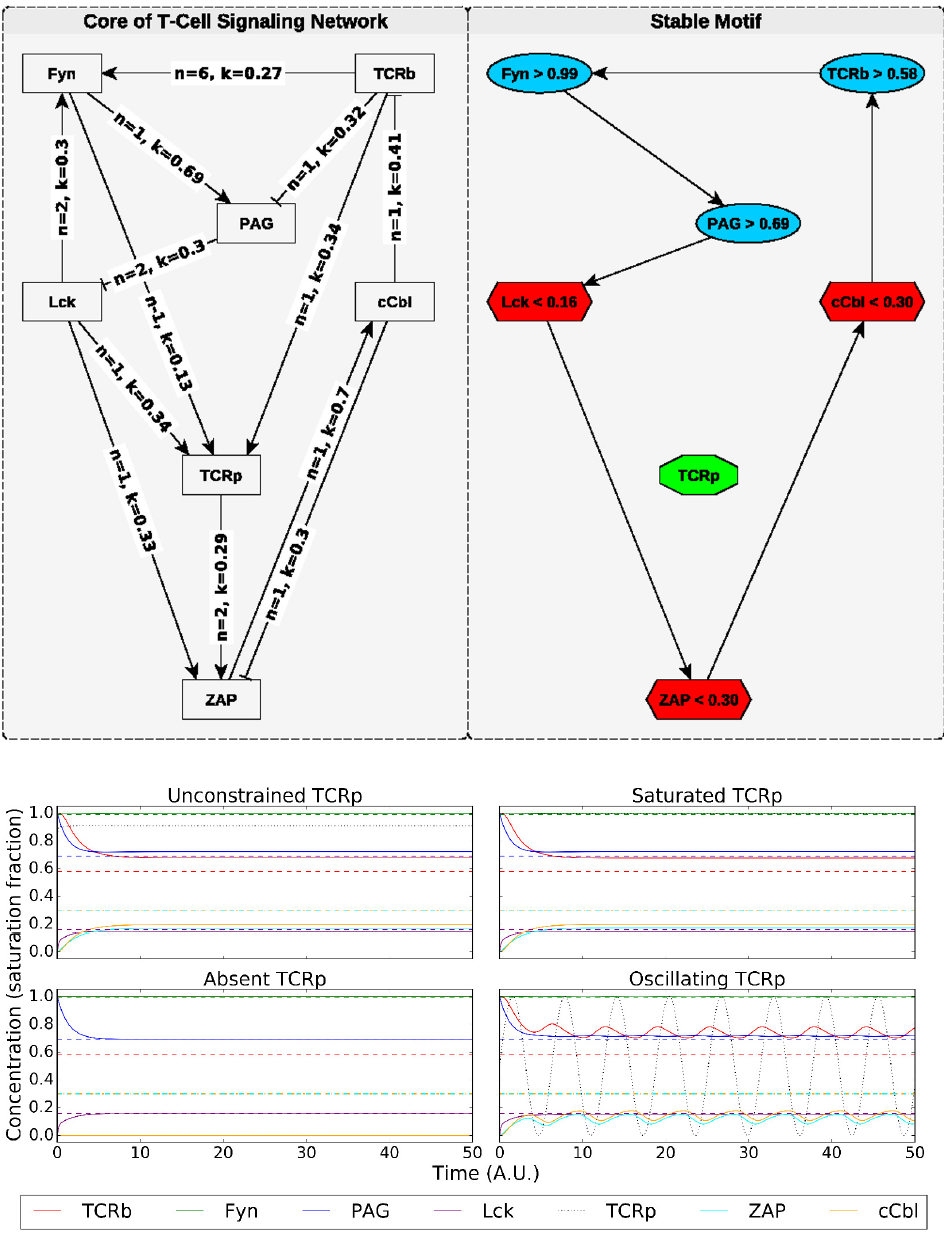
(a) The network diagram and stable motif for the T-cell signaling model of [37] with all sources saturated. In the stable motif diagram, node shape and color indicate whether an upper or lower bound is specified (as indicated by the node labels). The variables constrained by the stable motif cannot leave the region of state-space specified by the stable motif once it has been entered. This remains true even when *TCRp*, which regulates *ZAP*, but is not included in the stable motif, is subjected to external control, provided it remains within the bounds considered when constructing the expanded network (between 0 and 1 in this case, though a similar result can be obtained for 0 ≤*TCRp < ∞*). This robustness is illustrated in (b), in which solid colored lines indicate dynamic variable values that are constrained by stable motif thresholds (dashed lines). The black dotted line is the *TCRp* value and is subject to different external controls in panel of sub-figure (b).

By considering when the activation or inhibition of a given node is sufficient or necessary to cause the activation of other nodes, we have identified the cycle *TCRb →Fyn →PAG→ Lck →ZAP →cCbl →TCRb* as a candidate stable motif depicted in Fig 3 (right). This cycle is a positive feedback loop, but it is embedded in a sign-inconsistent network. As such, before we implement the MIOS approach to determine valid thresholds for the motif, we must address the effects of sign-inconsistent edges ([29]). For instance, in the motif, we expect *TCRb* and *PAG* to achieve relatively high values, but there is an inhibitory effect between the two; indeed,

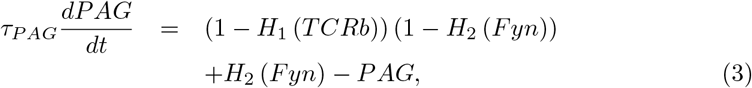

Where *H*_1_ and *H*_2_ are Hill functions (of the form 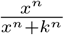 with *n* ∈ ℤ ^+^ and *k* ∈(0,1)). The inhibitory effect is maximized when *TCRb* attains its maximum value, i.e., one (because all variables are normalized to their maximum values in this model). It is also possible to consider the possibility that *TCRb* is delivered to the system via external control, in which case we would evaluate Eq. 3 in the limit as *TCRb→∞*. For now, we shall only consider *TCRb* = 1 in this regulatory function. We therefore replace *H*_1_ (*TCRb*) in Eq. 3 with *H*_1_ (1) and allow *TCRb* to evolve according to its natural dynamics in this new network, in which the regulation of *PAG* is modified. Similar analysis is taken on any edge that either introduces a sign inconsistency, or does not connect two nodes of the stable motif. The resulting modified network is a single positive feedback loop with a single steady state that is easily identified using the MIOS approach [29]. The steady state values of the nodes in the modified network serve as thresholds in the expanded network, and allow us to identify a stable motif (see Fig 3).

In this example, the stable motif we have identified coincides with a global steady state of the system. This observation is in agreement with [17], in which this system is analyzed by application of theorems regarding the conservation of certain positive invariant sets when a system is described by both a Boolean and an ODE model with Hill regulatory functions. We note that our analysis does not rely on a particular functional form of the regulation or on an explicit companion Boolean model. A novel result of our analysis is that the stable motif behavior cannot be disrupted by manipulating *TCRp*.

We demonstrate the robustness of the stable motif by numerically solving the system ODEs with various constraints placed on *TCRp* (Fig 3). In the top left panel of Fig 3, we show a natural evolution of the system for initial conditions satisfying the stable motif conditions. In the other panels, the value of the *TCRp* node is subjected to one of three external controls (absence, saturation, and oscillation), and the motif variables continue to respect the stable motif conditions. These simulations illustrate an important conclusion we can draw from the existence of the stable motif: If one wishes to avoid states in which *Fyn*, *PAG*, and *TCRb* are high while *Lck*, *ZAP*, and *cCbl* are low, *TCRp* is not a viable control target. Biologically, this model predicts that disruption of *TCR* phosphorylation is not sufficient to disrupt the response of the cell to a high degree of receptor-ligand binding. Instead one must disrupt one of the six motif nodes directly, and furthermore, the motif bounds provide lower bounds on the magnitude of the required disruption. For example, to disrupt the motif via control of *PAG*, one must lower its value below the threshold of 0.69.

## Discussion

We have presented a new framework, based upon construction of an auxiliary “expanded network”, for identifying self-sustaining subsystems that cannot be controlled via the rest of the system. Full attractor control requires that variables from each of these subsystems be externally manipulated. We have applied our framework to develop an algorithm (S1 Source Code) for finding these subsystems that is applicable in many biological ODE models. We have demonstrated our framework and algorithm in two biological systems: the T-cell receptor signaling network and the *Drosophila melanogaster* segment polarity gene network.

The method of expanded networks can extract important qualitative features from quantitative or qualitative models of system behavior. We have emphasized the identification of stable modules, which correspond to state-space regions that, once entered, cannot be exited without directly applying external control on the variables that define the region boundaries. We have also shown, for example in our analysis of the *Drosophila melanogaster* segment polarity gene network, how the consideration of expanded networks can elucidate meaningful and intuitive partitioning of state-space. In these analyses, we have considered virtual nodes of the form 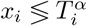, but other choices for virtual nodes are possible, and can be informative when *x_i_* has inherently multi-level behavior.

In searching for stable modules, it is important to identify positive feedback loops, as every stable module of the type considered in our automation procedure corresponds to a sign-consistent subgraph that must contain at least one positive cycle. In the examples described here, as well as in every ODE model of biological systems we encountered so far, the number of positive feedback loops is small, and so an exhaustive search is feasible. Even when this number is large, an exhaustive test of all sign-consistent subgraphs is probably still faster than a brute-force simulation approach because this method is testing many control strategies simultaneously for each subsystem, and does not involve integration of any ODEs. Positive feedback loops can be identified using existing software implementations. For each positive feedback loop, the computational complexity of determining the associated thresholds scales linearly with the number of variables in the feedback loop. If necessary, we are also able to limit our search to positive feedback loops of a certain size, allowing for fast identification of small, control-robust subsystems embedded in much larger systems.

Our procedure yields all stable modules of threshold statements about variables involved in positive feedback loops. These correspond to positive-invariant sets that remain positively invariant even when regulatory effects external to the feedback loop are manipulated. Because we cannot know *a priori* which regulatory manipulations are available within a given model, we have chosen to focus on behaviors that are robust to *all* manipulations of these external regulations. Therefore, the stable modules we identify are robust to control beyond that which can be implemented in practice. In some systems, additional behaviors may exist that are robust only to a biologically relevant subset of control strategies. Nevertheless, knowledge of the fully robust system behaviors reduces (in some cases, dramatically) the search space for control targets. For example, in a large system, we might identify a stable module that includes some “undesirable” behavior (e.g., a disease state) and involves a small number of variables. Because the stable module subsystem is robust to all topology-preserving external controls, control targets must be selected from the small number of variables directly involved in the stable module.

Many existing results about the analysis of Boolean models via expanded networks remain valid in this more general framework and can therefore be applied to continuous systems. An example is the concept of a driver node set, which is a set of virtual nodes in the expanded network whose truth eventually implies the truth of a given stable module. Identification of driver nodes in the expanded network is related to finding paths in logic hypergraphs [31]. This identification problem has been partially addressed in [30, 39]; developing a general and fast algorithm for driver node identification in arbitrary expanded networks is a promising direction for future research with applications for control target selection.

Some results do not generalize as easily because they rely upon completeness properties of discrete expanded networks; the oscillation analyses in [30, 33] are an example. Oscillations can manifest in the expanded network as source-free graph components that contain contradictory nodes. Such structures do not always indicate oscillatory behavior, and may instead indicate chaotic behavior or the existence of a steady state that violates all of the contradictory conditions. For example, the simple harmonic oscillator *x*̇ = *y*, *y*̇ =–*x* has an expanded network with contradictory source-free component *x >* 0 →*y <* 0 →*x <* 0 →*y >* 0; while the system can oscillate between satisfying these conditions, there is also a steady state, *x* = *y* = 0, that violates all four conditions. We are optimistic that results of this type might be recast in more general forms.

The expanded network framework shows promise not only for studying the state-space of dynamical systems, as we have emphasized here, but also for the study of parameter space. Statements regarding the value of parameters can be included in an expanded network as statements with self-loops. Because the expanded network approach extracts qualitative information from the system, the inclusion of parameters in this way is conceptually distinct from and complementary to existing methods for probing the parameter space of a dynamical system (e.g., [40, 41]). The application of expanded networks to parameter sensitivity analyses is the subject of ongoing work.

## Supporting information

**S1 Appendix. Supplementary Notes and Examples.**

**S1 Source Code. Six python source files that contain the implementation of the stable module search procedure and its application to five examples used in the main text and S1 Appendix.**

**S1 Figure. Results of numerical simulations of the hypothetical**

***Drosophila* cell transplant experiment discussed in the main text.**

## Acknowledgments

We thank Jorge Zañudo for his helpful insights and conversations.

